# Stress-responsive *Entamoeba* topoisomerase II: a potential anti-amoebic target

**DOI:** 10.1101/679118

**Authors:** Sneha Susan Varghese, Sudip Kumar Ghosh

## Abstract

Topoisomerases are ubiquitous enzymes, involved in all DNA processes across the biological world. These enzymes are also targets for various anticancer and antimicrobial agents. The causative organism of amoebiasis, *Entamoeba histolytica* (*Eh*), has seven unexplored genes annotated as putative topoisomerases. One of the seven topoisomerases in this parasite was found to be highly up-regulated during heat shock and oxidative stress. The bioinformatic analysis shows that it is a eukaryotic type IIA topoisomerase. Its ortholog was also highly up-regulated during the late hours of encystation in *E. invadens* (*Ei*), the encystation model of *Eh*. Immunoprecipitated endogenous EhTopoII showed topoisomerase II activity *in vitro*. Immunolocalization studies show that this enzyme colocalized with newly forming nuclei during encystation, which is a significant event in maturing cysts. Double-stranded RNA mediated down-regulation of the TopoII both in *Eh* and *Ei* reduced the viability of actively growing trophozoites and also reduced the encystation efficiency in *Ei*. Drugs, targeting eukaryotic topoisomerase II, e.g., etoposide, ICRF193, and amsacrine, show 3-5 times higher EC_50_ in *Eh* than that of mammalian cells. Sequence comparison with human TopoIIα showed that key amino acid residues involved in the interactions with etoposide and ICRF193 are different in *Entamoeba* TopoII. Interestingly, ciprofloxacin an inhibitor of prokaryotic DNA gyrase showed about six times less EC_50_ value in *Eh* than that of human cells. The parasite’s notable susceptibility to prokaryotic topoisomerase drugs in comparison to human cells opens up the scope to study this invaluable enzyme in the light of an antiamoebic target.

## Introduction

Amoebiasis is a leading cause of death due to a parasitic protozoan infection worldwide, highly prevalent in regions with poor sanitation and hygiene. The causative organism, *Entamoeba histolytica*, alternates between two life stages- pathogenic trophozoite and resistant cyst stage [1]. Stage conversion to the dormant form is called encystation and is the critical process for host-to-host transmission of the disease. Much of the information about encystation comes from studying the closely related reptilian parasite *E. invadens*, as it can encyst *in vitro* under reduced osmotic pressure and nutrient depletion [2, 3]. Hence, stress is a precursor for encystation, and this process witnesses the activation of several specific stress-responsive proteins and signaling pathways [4, 5]. One of the key features during this stage conversion is the appearance of four distinct nuclei during the later hours of encystation [6, 7]. Association of the sexual process, like meiosis, with encystation in *Entamoeba*, has been a point of debate for long. Low levels of allelic heterozygosities [8], evidence of homologous recombination [9], the presence of meiotic genes and their upregulation during encystation [10], and the formation and aggregation of haploid nuclei in multinucleated giant cells during encystation [11] indicate the occurrence of meiosis-like processes and homologous recombination during glucose deprived encystation. Across the biological world, all key DNA processes are associated with a group of ubiquitous enzymes called topoisomerases. These enzymes maintain DNA topology by creating breaks in the DNA and allowing strand passage. Thus, these are involved in removing DNA supercoils, chromosomal condensation, strand-breakage during recombination, and disentangling intertwined DNA, in naming a few [12, 13]. Based on their mode of action, there are two broad types of topoisomerase, viz Type I and Type II. The former function as monomers, creating single strand breaks while the latter forms multimers and introduces double-strand breaks in the DNA. Owing to the vital functions of these enzymes, they have been extensively evaluated as targets for many antitumors, antibacterial, and antifungal drugs [14–17]. Also, topoisomerases in several pathogenic protozoans like *Plasmodium*, and *Leishmania* are being evaluated as targets for antiparasitic agents [18–21].

Treatment for amoebiasis relies primarily on nitroimidazole based drugs, especially metronidazole [22]. Although effective both in bowel lumen and tissue, this broad spectrum drug is reported to eradicate only 50% of luminal infection [23]. Successful induction of metronidazole-resistant parasitic strain at the laboratory level points to the probability of drug resistance in the near future [24, 25]. Moreover, the side effects of metronidazole include nausea, abdominal pain, diarrhea, and in severe cases, results in neurotoxicity, optic neuropathy, peripheral neuropathy, and encephalopathy [26, 27]. Hence, there is a need for newer and effective alternative solutions.

In this study, stress-responsive, eukaryotic Type IIA topoisomerase was identified in *Eh* and *Ei*, upregulated both at RNA as well as protein level during later stages of encystation, and various other stresses, like heat shock and oxidative stress. RNAi mediated silencing of that gene has shown a significant impact on the overall viability and encystation efficiency of *Entamoeba*. Further, a higher potency of prokaryotic topoisomerase II drugs on the parasite when compared to human cells makes it a significant drug target worth further exploration.

## 2. Results

### 2.1. Phylogenetic classification of *Entamoeba* topoisomerase genes

Topoisomerases are broadly categorized as type I and II, based on their structure and function. Type I functions as a monomer, creating single strand breaks and based on differences in the mechanism of action, they are subcategorized as IA and IB. Prokaryotic topo I, all topo III and reverse gyrase belong to the former while latter comprises of eukaryotic topo I and archeal topoV. Although all type II topoisomerases are multimers and create double strand breaks, based on their structural differences, these are sub-classified as type IIA (eukaryotic topo II, topo IV and DNA gyrase) and type IIB (topo VI).

Phylogenetic analysis shows that all genes annotated as topoisomerases in *Entamoeba* are eukaryotic (Fig 1). Interestingly, orthologs of *Eh* and *Ei* were very closely related, and hence observations for one may be extrapolated to its ortholog. EHI_073170/EIN_344850 and EHI_087330/EIN_173080 showed a very distant relation with Topo VI, which is reported only in plants and archaeabacteria. From the remaining pool of five genes, EHI_038920/EIN_052260 and EHI_042880/EIN_174490 shared maximum similarity with topoisomerase IIIα and IIIβ (Type IA), respectively. Further, EHI_125320/EIN_371990 and EHI_194510/EIN_229710 aligned closely with SPO11 family, which function similar to Topoisomerase II and creates double-strand breaks during meiosis. EHI_120640/EIN_145900 resembled eukaryotic Topoisomerase IIα (Type IIA). Multiple sequence alignment and NCBI-CDD shows that the putative topoisomerase II is a dimer with three core domains: N-terminal ATPase domain, a central domain carrying the active site and a C-terminal variable domain carrying the NLS sequence (Fig 2). Interestingly, neither of the *Entamoeba* species had a recognizable eukaryotic topo I (Type IB).

**Fig 1.**
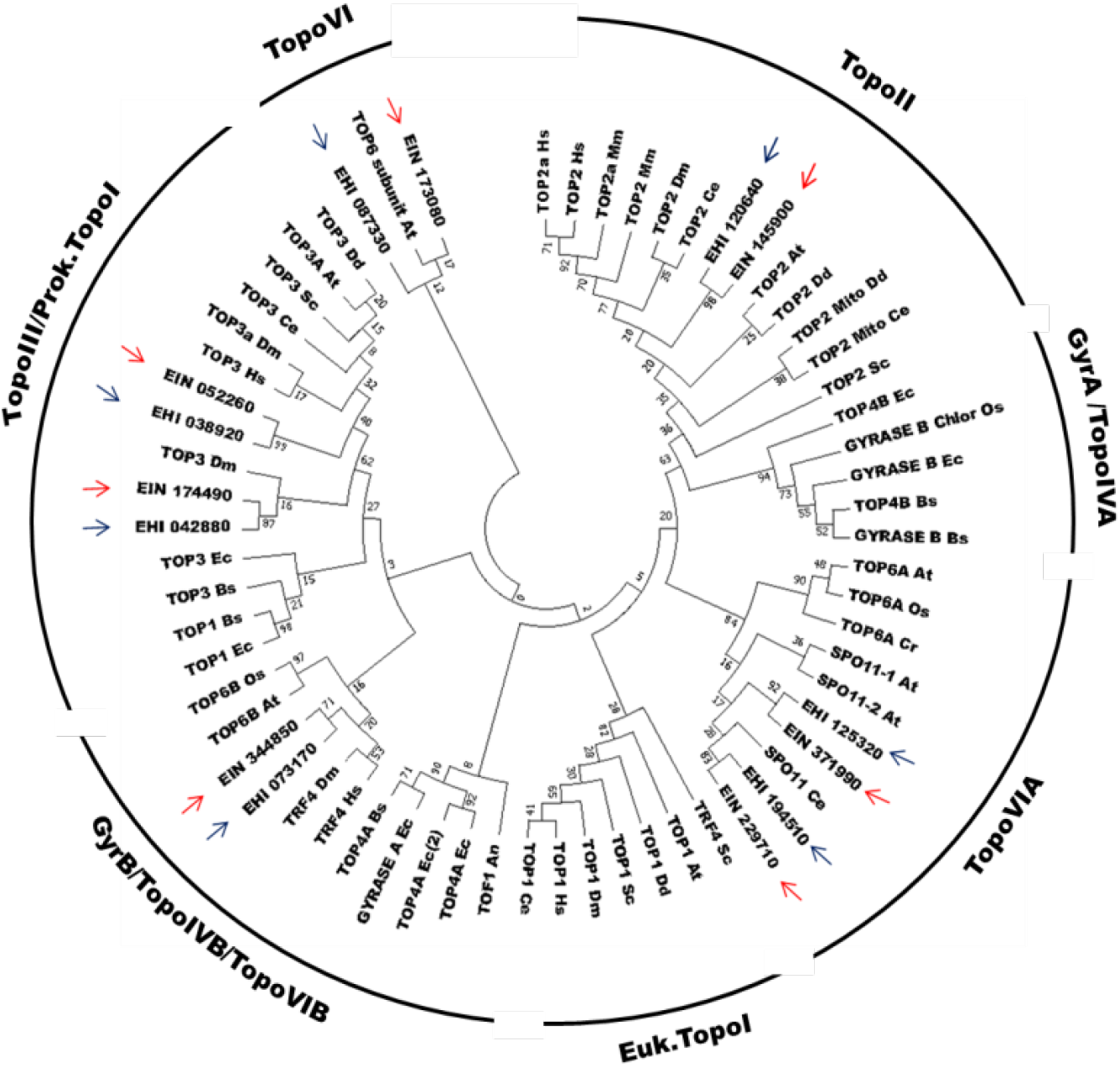
The phylogenetic classification of *Entamoeba* topoisomerase genes. Phylogenetic tree generated based on maximum likelihood method using MEGA 7.0 software. Different topoisomerases from various prokaryotic and eukaryotic organisms were used to construct the tree. The seven putative topoisomerases of *E. histolytica* and *E. invadens* is highlighted with blue and red arrows, respectively. **At:** *Arabidopsis thaliana*; **Bs**: *Bacillus subtilis*; **Ce**: *Caenorhabditis elegans*; **Cr**: *Chlamydomonas reinhardtii*; **Ds**: *Dictyostelium discoideum*; **Dm**: *Drosophila melanogaster*; **Ec**: *Escherichia coli*; **Hs**: *Homo sapien* **Mm**: *Mus musculus*; **Os**: *Oryza sativa*; **Sc**: *Saccharomyces cerevisiae*.

**Fig 2.**
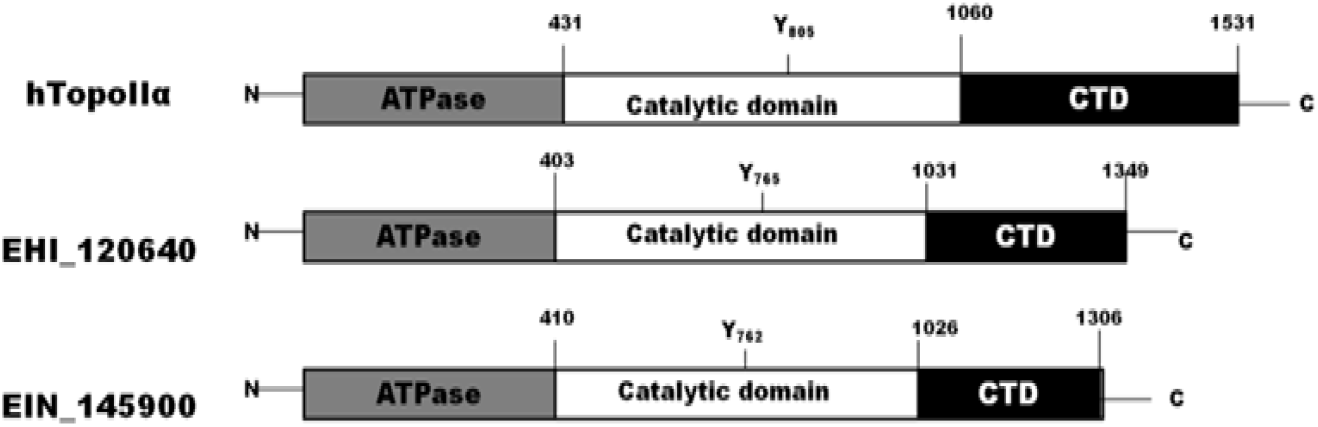
Domain organization on putative Topo II of *Entamoeba*. Comparison of predicted domains on putative topoisomerase II of *Eh* and *Ei* with the domains of human topoisomerase IIα. The putative TopoII of *Entamoeba* has all the three crucial domains: N-terminal domain with ATPase activity, Central catalytic domain housing the active Tyr residue, and highly variable C-terminal domain that nestles the Nuclear Localization Sequence.

### 2.2. Expression profile of putative *Entamoeba* Topoisomerase genes during different stresses and encystation

Real-time RT-PCR analysis of these five genes shows that only EHI_120640 and its ortholog EIN_145900 were highly upregulated during heat shock and oxidative stress, although the gene expression did not significantly alter during 16h of glucose starvation. As a similar upregulation of EIN_145900 gene expression was also observed during later periods of encystation, this could be important for stress response in *Entamoeba* (Fig 3). Much like during 16h of glucose starvation, early periods of encystation did not show upregulation of EIN_145900. Similar observations for all these genes during encystation were noted from the microarray data [28] (S1 Fig). As a result, further study focused on this stress responsive putative topoisomerase II. EIN_229710, predicted as SPO11 has already been reported to be a meiosis-specific gene and upregulated during encystation of *E. invadens* [10].

**Fig 3.**
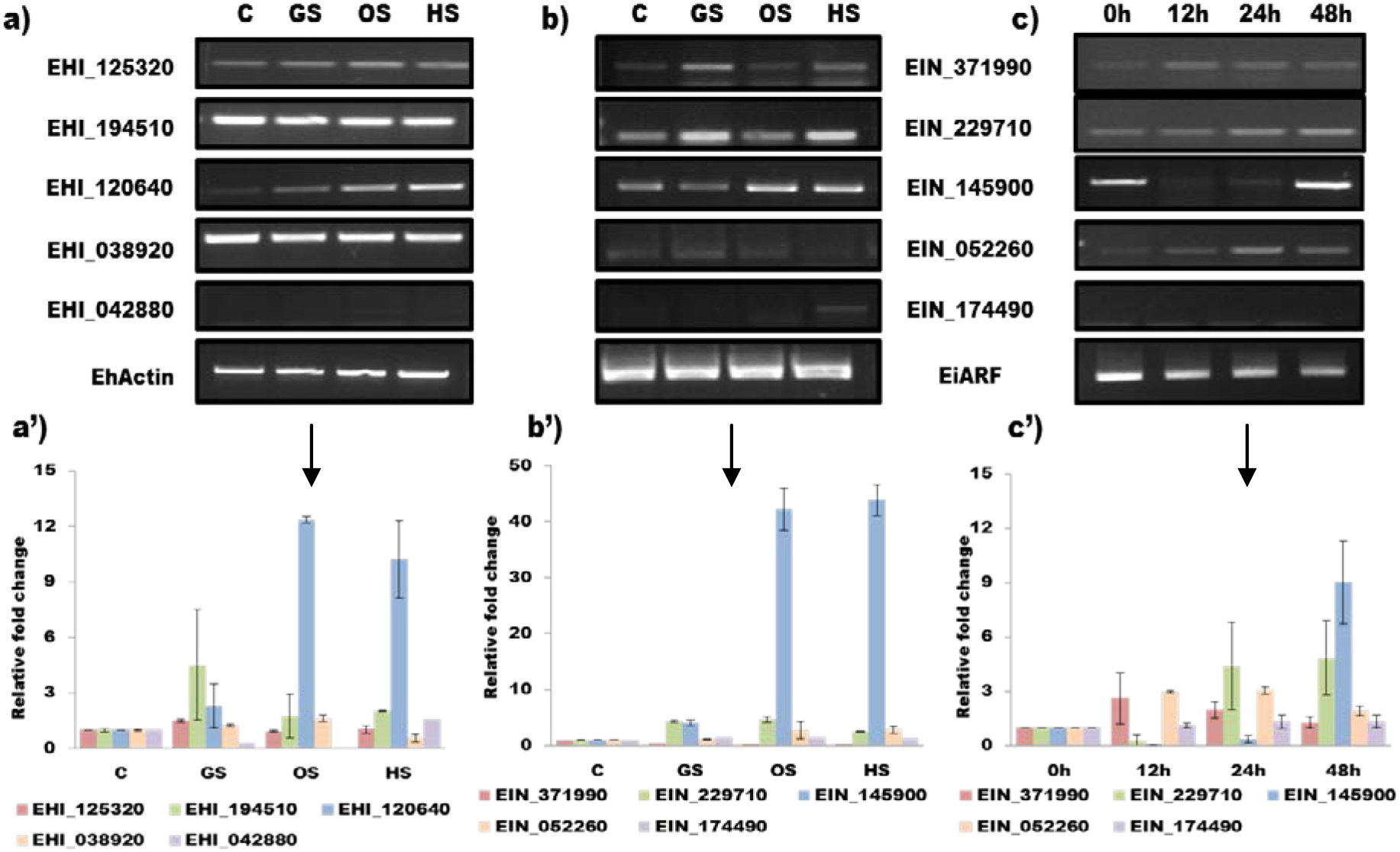
Expression profile of putative *Entamoeba* Topoisomerase genes during different stresses and encystation. sq-RT PCR and corresponding Real-time PCR analysis showing relative fold change in transcription of different putative topoisomerases under different stress conditions of **a)** *E. histolytica* and **b)** *E. invadens* and, **c)** *E. invadens* during different hours on encystation. EHI_120640 and its ortholog, EIN_145900 are markedly upregulated during various stress and later hours of encystation. **C**-Control; **GS**-Glucose starvation; **OS-** Oxidative stress; **HS-** Heat shock

### 2.3. Cloning, expression, and purification of recombinant putative EhTopo II fragment

Although different combinations of bacterial expression vector and strains were tried, full-length expression of putative EhTopo II (4050bp) was not successful. Hence a 1127bp fragment (from 760bp-1887bp) was successfully cloned into pET21a expression vector, and recombinant EhTopo II fragment was successfully expressed with a 6x Histidine tag at the C-terminal in *E. coli* Bl21(DE3) strain using 1mM IPTG. The insoluble, recombinant protein so obtained was solubilized using 0.25% S-lauryl sarcosine, purified by Ni-NTA affinity chromatography and resolved by SDS-PAGE (S2 Fig). Purified recombinant EhTopo II fragment was used to raise antibody in rabbits. Anti-EhTopoII antibody was purified from crude sera and confirmed by Western blot analysis. As the antigenic fragment of EhTopo II shared 70% sequence similarity with its ortholog in *Ei*, the anti-EhTopo II antibody could successfully bind to EiTopo II as well (S3 Fig).

### 2.4. Topoisomerase II protein expression is upregulated during different stresses and encystation in *Entamoeba*

Topoisomerase II protein was expressed under all the studied stress conditions in both *E. invadens* and *E. histolytica* with a marked upregulation during heat shock and oxidative stress (*p<0.05, as compared to control) (Fig 4a and 4b). The temporal expression of the protein during encystation followed a pattern similar to the transcript profile wherein the protein expression gradually declined in the early hours and showed a drastic upregulation during the later stages, which is also the period of tetranuclei formation (Fig 4c). Hence this confirms *Entamoeba* Topo II is upregulated at both transcription and protein synthesis.

**Fig 4.**
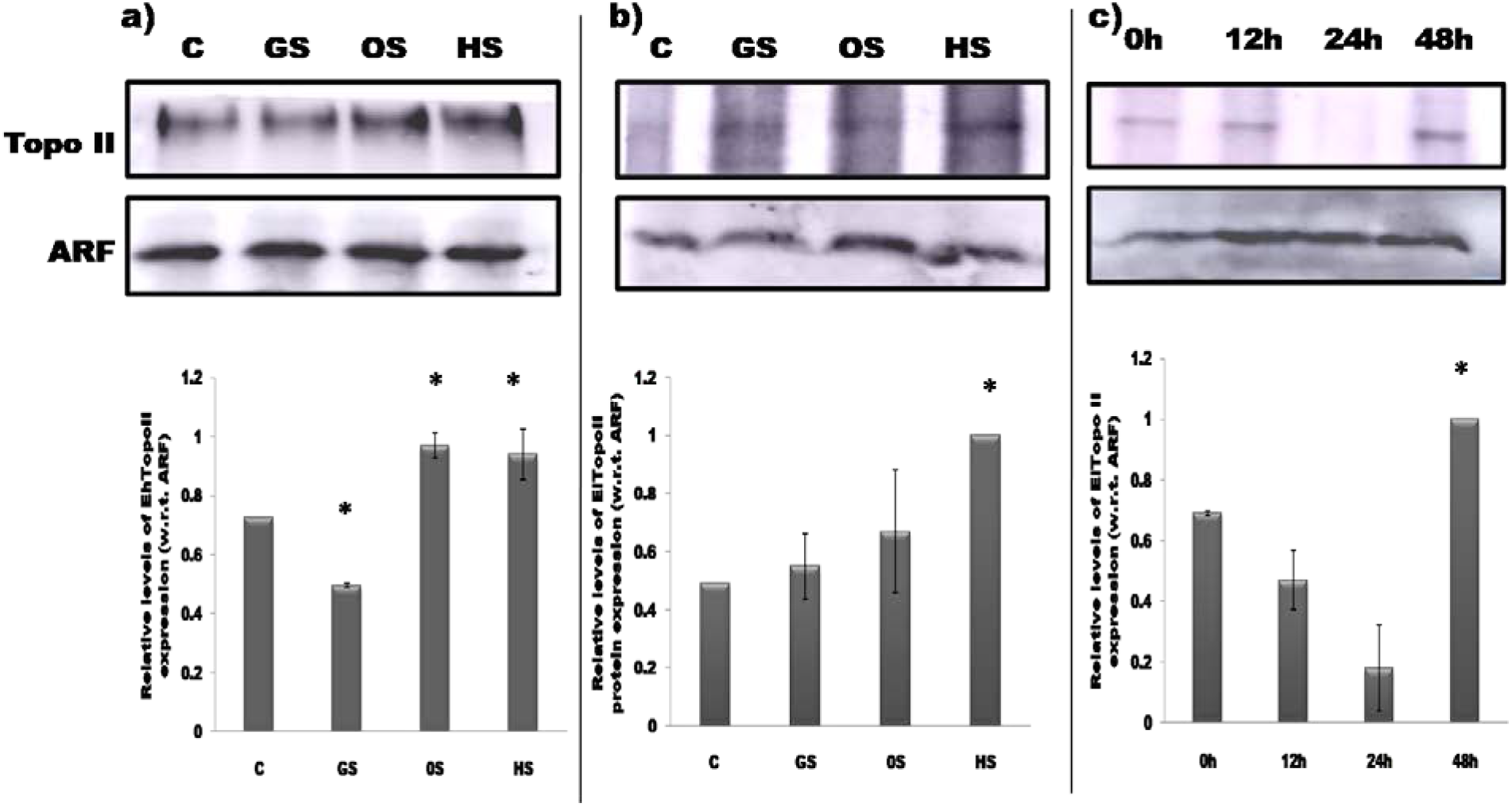
Upregulation of Topoisomerase II protein expression during different stress and encystation in *Entamoeba*. Western blot analysis for the expression profile of Topo II protein during different stresses in **a**) *E. histolytica*, **b**) *E. invadens* and **c**) during different hours of encystation in *E. invadens*, respectively. Similar findings as that of Real-time RT-PCR were observed. ARF was used internal control for normalisation. **C-** Untreated control; **GS-** Glucose starvation; **OS-**Oxidative stress; **HS-** Heat shock.

### 2.5. *Entamoeba* trophozoites express functional topoisomerase II

The function of topoisomerases is to relieve the topological strain that occurs in the DNA during various processes like replication, transcription, chromosomal segregation, etc. This is achieved by relaxing positive or negative supercoils in the DNA and depending on the type of enzyme, various cofactors are required for this process. We have predicted the presence of eukaryotic Topo III and Topo II in *Entamoeba*. TopoIII requires only MgCl_2_ while Topo II requires both ATP and MgCl_2_ as cofactors for its enzymatic activity. To determine whether *Entamoeba* trophozoites express functional topoisomerase II, we studied its ability to relax negative supercoils. Varying concentration of nuclear extracts, prepared from actively proliferating cultures, could successfully relax negatively supercoiled plasmids in the presence of co-factors ATP and MgCl_2_. At 40μg of crude nuclear extract, *Eh* was able to relax 100% of the supercoiled plasmid in comparison to 70-80% relaxation by *Ei* at the same concentration (Fig 5a and 5b). This study shows the presence of active Topo II in *Entamoeba* trophozoites

**Fig 5.**
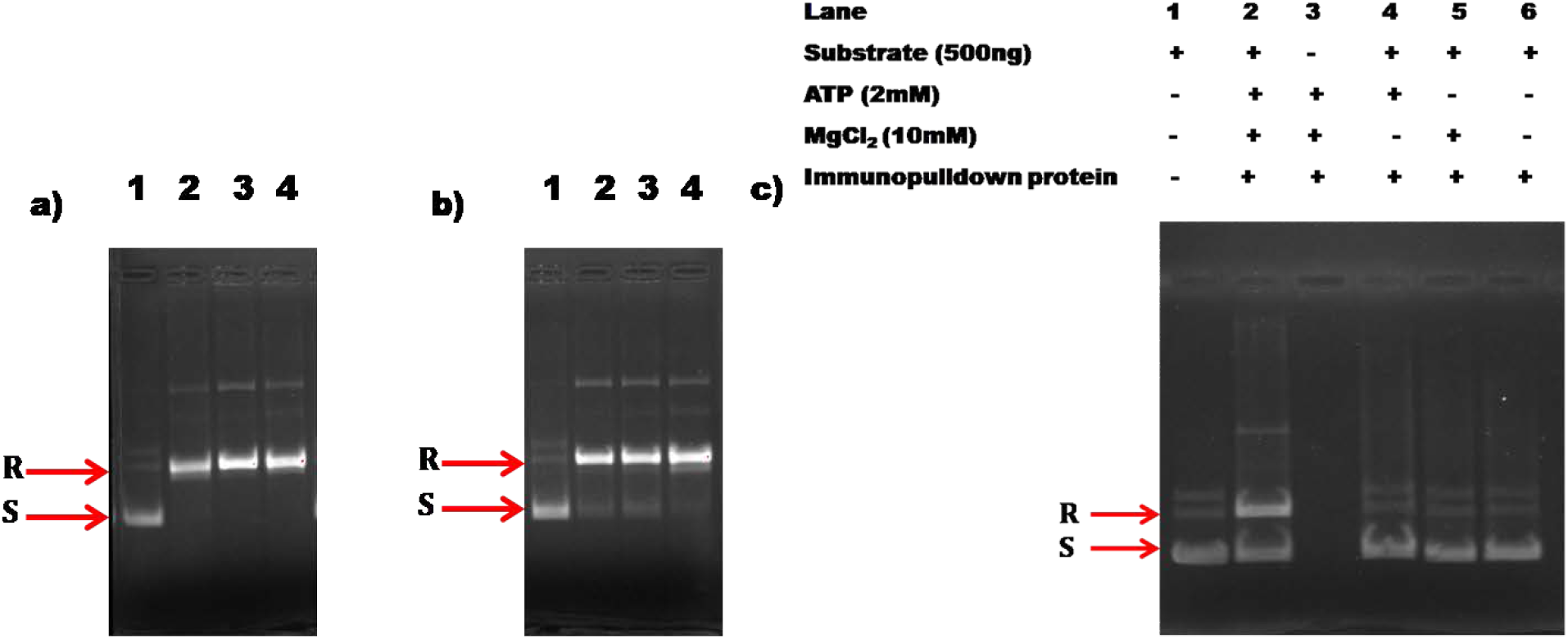
*Entamoeba* trophozoites express functional topoisomerase II. Agarose gel electrophoresis (1%) depicting the relaxation of negatively supercoiled plasmids by varying amounts of crude nuclear (Lane 1-4 contains 0, 20μg, 40μg and 60μg of crude nuclear extract from the respective source) of **a)** *Eh* and **b)** *Ei*. **c)** endogenous EhTopoII (immunoprecipitated from nuclear extracts of *Eh*, using anti-TopoII antibody). Relaxation activity occurs only in the presence of both ATP and MgCl_2_. **R** and **S** represent the negatively supercoiled plasmid and the relaxed form, respectively.

Endogenous EhTopo II was immunoprecipitated from the crude nuclear extract using anti-EhTopo II antibody. The native EhTopo II showed relaxation activity only in the presence of both ATP and MgCl_2_. As no relaxation was observed in the absence of either ATP or MgCl_2_ or both, it can be concluded that the pulled down Topo II was not contaminated with any type I topoisomerases (Fig 5c).

### 2.6. Localization of Topoisomerase II on newly forming tetranuclei during encystation in *Entamoeba*

Confocal micrograph analysis showed that Topo II localizes in the nucleus of *Eh* trophozoites during normal growth as well as during oxidative stress and heat shock. Contrarily, during glucose starvation, the protein was observed to move out into the cytoplasm (Fig 6a), and this observation during glucose starvation was consistent in *Ei* as well (Fig 6b). During the early hours of encystation (12 hours) EiTopo II expression reduced in comparison to trophozoite. This observation is in consensus with real-time RT-PCR and western blot analysis. As topoisomerase II is a key player in cell growth and proliferation, degradation of the enzyme during glucose starvation and early encystation could be a part of the initial survival response of the organism to energy deficiency. However, with the progression of encystation into later hours, Topoisomerase II co-localized with the newly forming tetranuclei suggesting that this enzyme may be responsible for relieving the topological strains that occur in the DNA during this nuclear event (Fig 6c).

**Fig 6.**
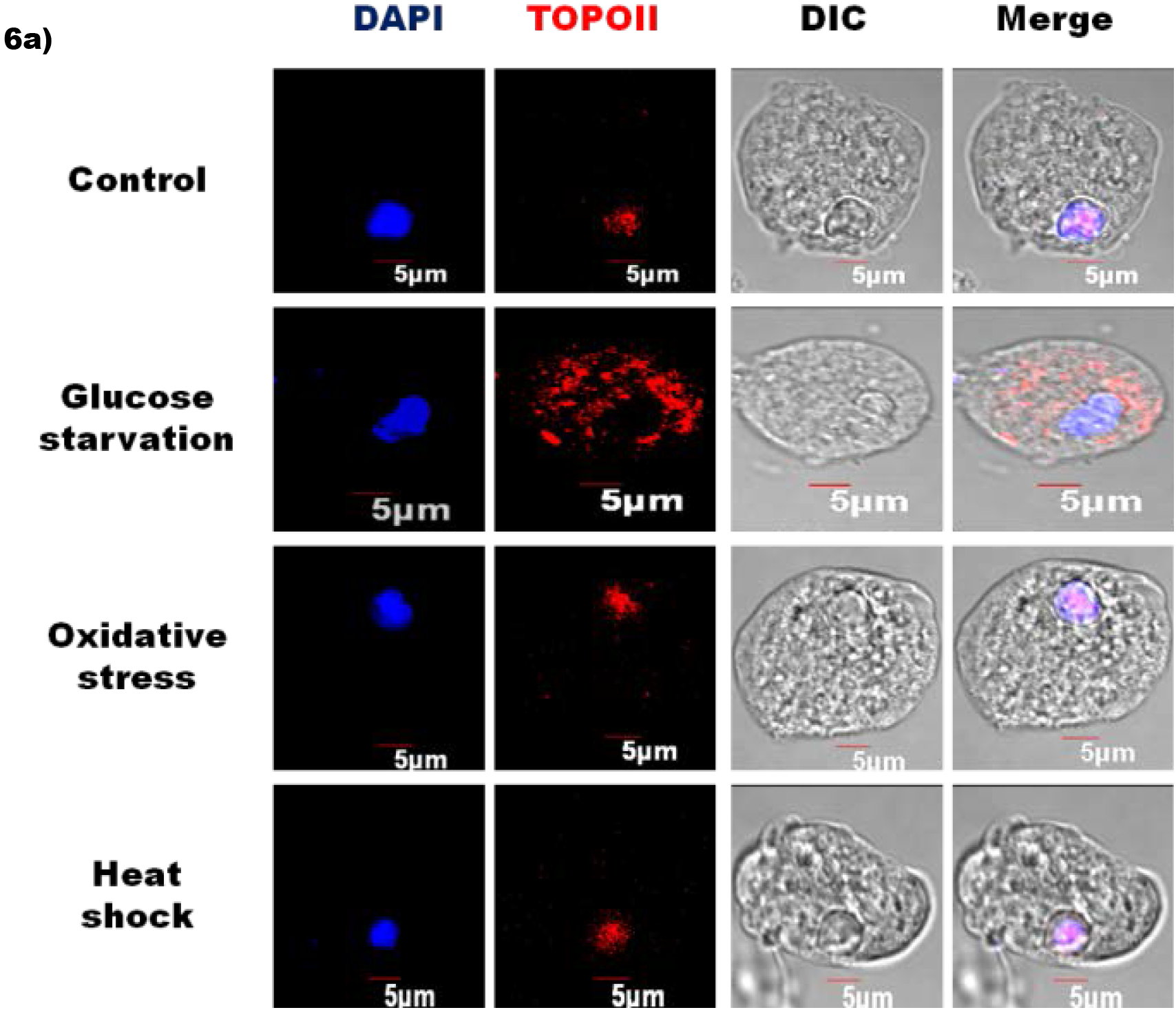
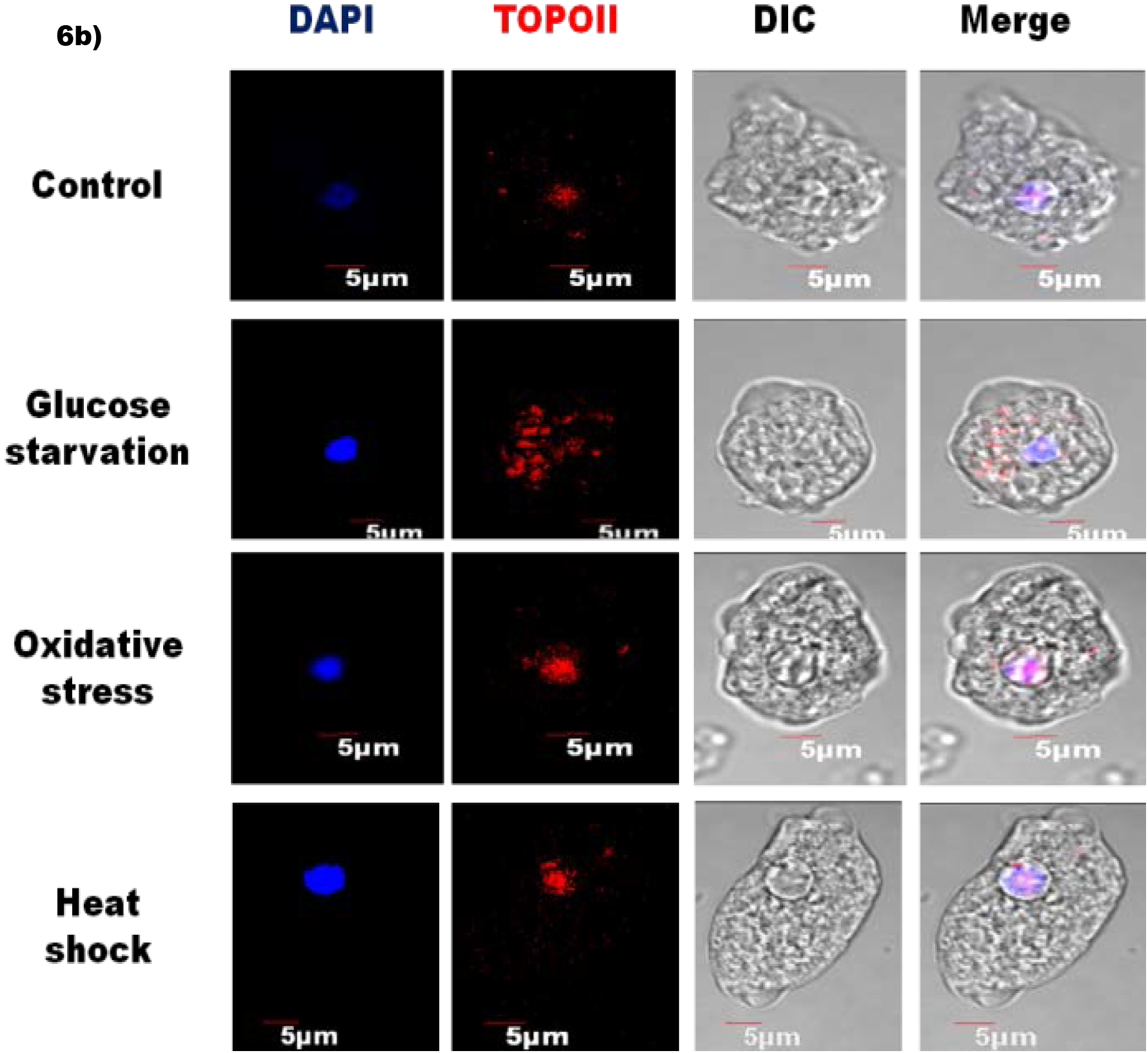
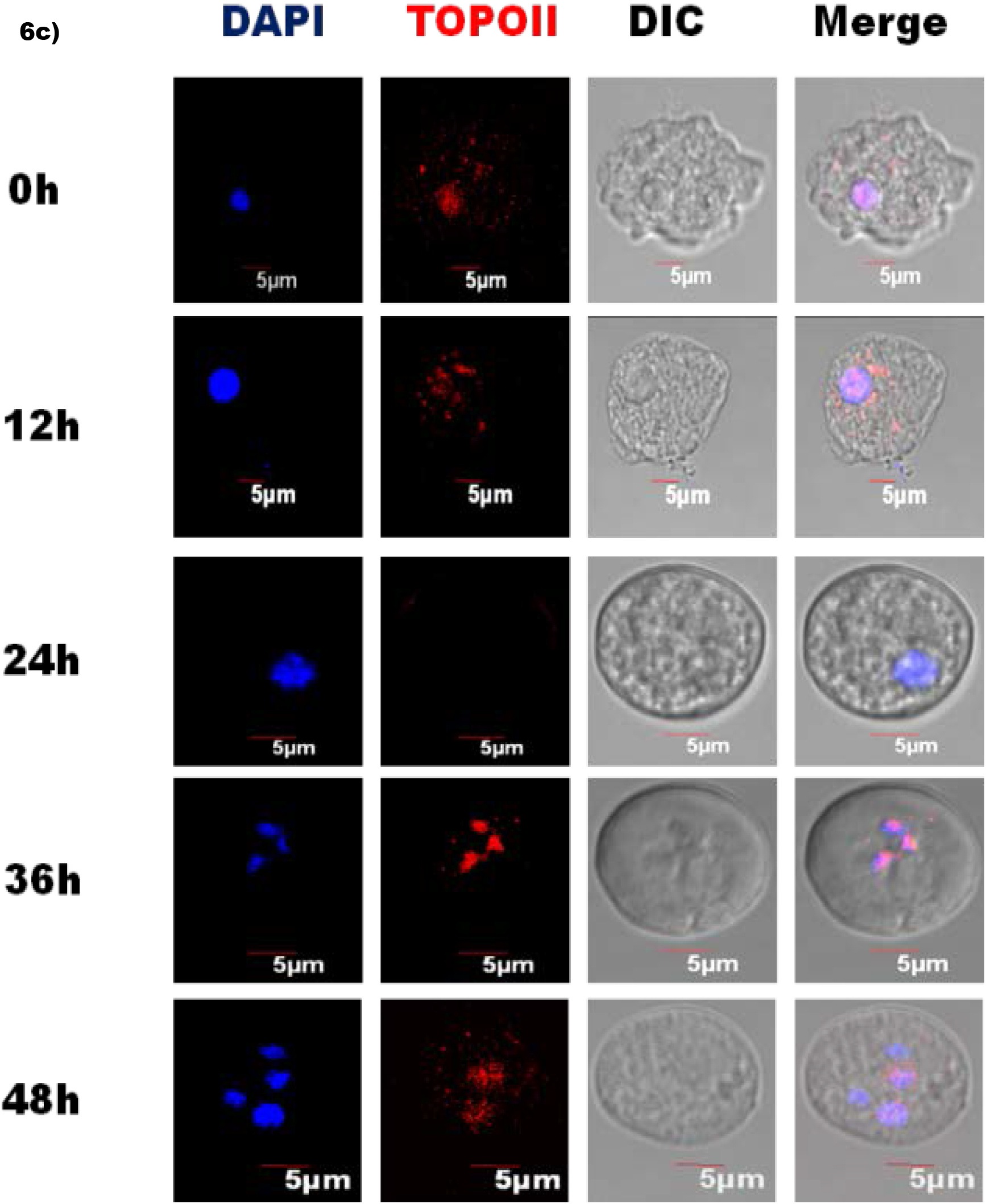
Topo II localization during stress and encystation in *Entamoeba*. The localization pattern of Topo II during **a)** different stress conditions in *Eh* and **b)** *Ei*. The enzyme primarily localized in the nucleus, except during glucose starvation. **c)** Localization of Topo II during different hours of encystation in *E. invadens*. The early hours of encystation show the decline of topoisomerase II from the cell. However, during the later stages of encystation, the protein reappears and localizes on the newly forming nuclei. *Entamoeba* TopoII is stained red with TRITC, and the nucleus is stained blue with DAPI.

### 2.5. TopoII silencing in *Entamoeba* significantly affects the viability of trophozoite and cyst

To understand the *in vivo* role of *Entamoeba* TopoII during normal growth as well as encystation, we employed the dsRNA mediated silencing strategy reported by Samanta and Ghosh, 2012 [29]. Gene-specific dsRNA was cloned into the pL4440 vector which is flanked by T7 promoter on either side, expressed and purified from RNase III-deficient *E. coli* HT115 cells. Approximately, 60-70% reduction in the transcription of Entamoeba *Topo II* was observed in samples soaked with 200μg/mL gene-specific dsRNA in comparison to control (untreated) (S4 Fig).

Topoisomerase is considered a proliferation marker and is an absolute requirement for rapidly proliferating eukaryotic cells. Interestingly, silencing of topoisomerase II in actively growing, log-phase trophozoites significantly (***P<0.001) reduces the viability by about 25% and 30% in *Eh* and *Ei*, respectively (Fig 7a). A similar reduction in viability as well as encystation efficiency was observed upon silencing the gene in samples under encystation condition. The encystation efficiency and viability of *EiTopoII* silenced cells reduced by approximately 42% and 28% respectively in comparison to the two control conditions (Fig 7b and 7c).

**Fig 7.**
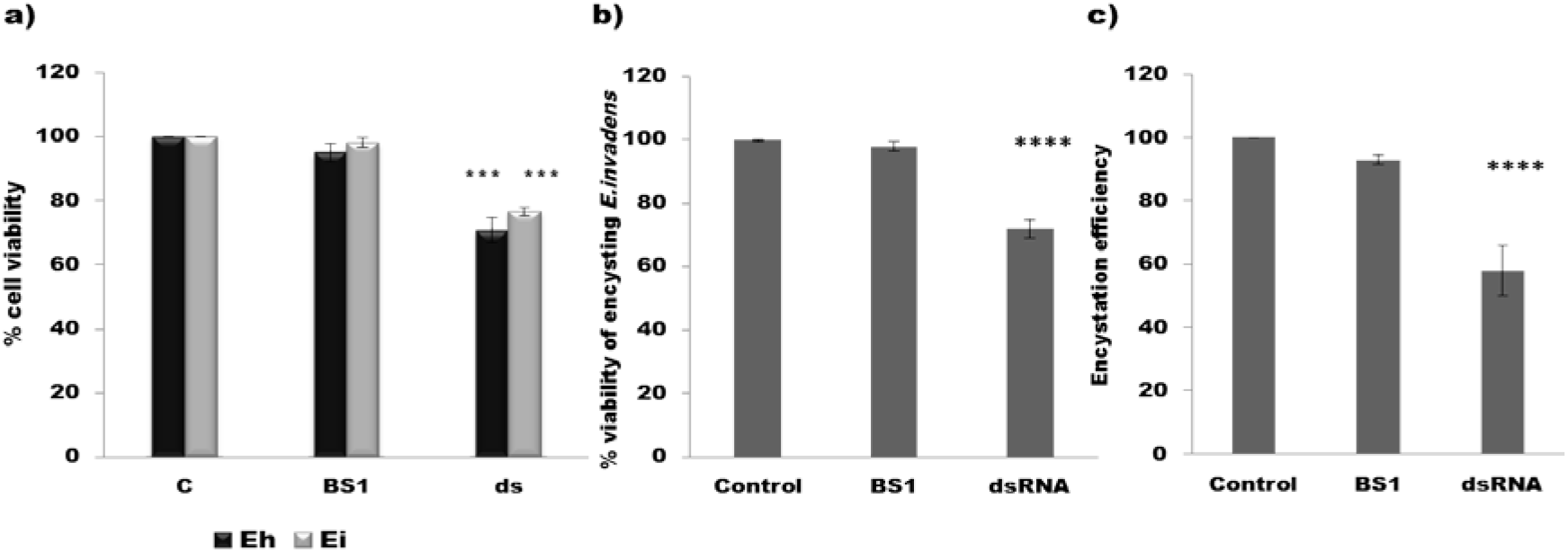
TopoII silencing in *Entamoeba* significantly affects the viability of trophozoite and cyst. **a)** The % viability of *Eh* and *Ei* trophozoites, on silencing the respective Topoisomerase II for 24h. A significant reduction of 30% and 25% (P<0.001) respectively, in the viability of actively growing trophozoites, were observed. **b)** Cell viability (%) and **c)** encystation efficiency of *Ei* after 48h of encystation during dsRNA mediated Topo II downregulation (200μg/ml). A significant reduction (P<0.0001) in the viability and encystation efficiency of about 30% and 42% respectively observed upon silencing *TopoII*

### 2.6. Less toxicity of eukaryotic topoisomerase II inhibitors in *E. histolytica*

As topoisomerase II is an extensively studied class of enzyme and is essential for cell growth and proliferation, many drugs designed against the eukaryotic topoisomerase II are available as anticancer and antifungal agents. We tested the potency of three different kinds of eukaryotic Topoisomerase II inhibitors viz. etoposide, amsacrine, and ICRF-193 on *E. histolytica*. The former two stabilize the cleavable complex, prevent the religation of double-strand breaks and hence stimulate enzyme-mediated DNA breakage. ICRF 193, a bizdioxopiperazine, locks the enzyme in a closed-clamp and prevents ATP hydrolysis necessary to regenerate the active form of the enzyme. Etoposide, ICRF-193, and amsacrine were lethal against *E. histolytica* with EC_50_ of 200μM, 15μM and, 80μM, respectively (Fig 8). However, the susceptibility of *Eh* to eukaryotic drugs is approximately 3-5 times lower than that of the human hosts, especially towards etoposide (Table 1).

**Fig 8.**
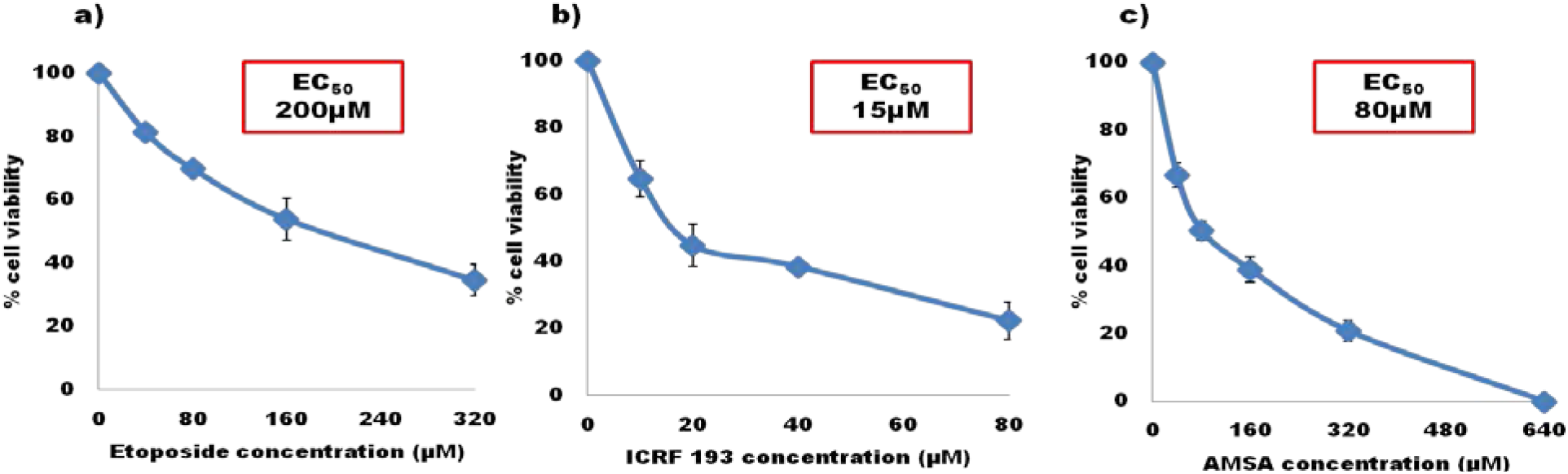
Less toxicity of eukaryotic topoisomerase II inhibitors in *E. histolytica*. Dose-dependent effect of eukaryotic Topoisomerase II drugs, namely **a)** etoposide **b)** ICRF 193 and **c)** amsacrine (AMSA) on *E. histolytica* viability following 24h treatment.

**Table 1.**
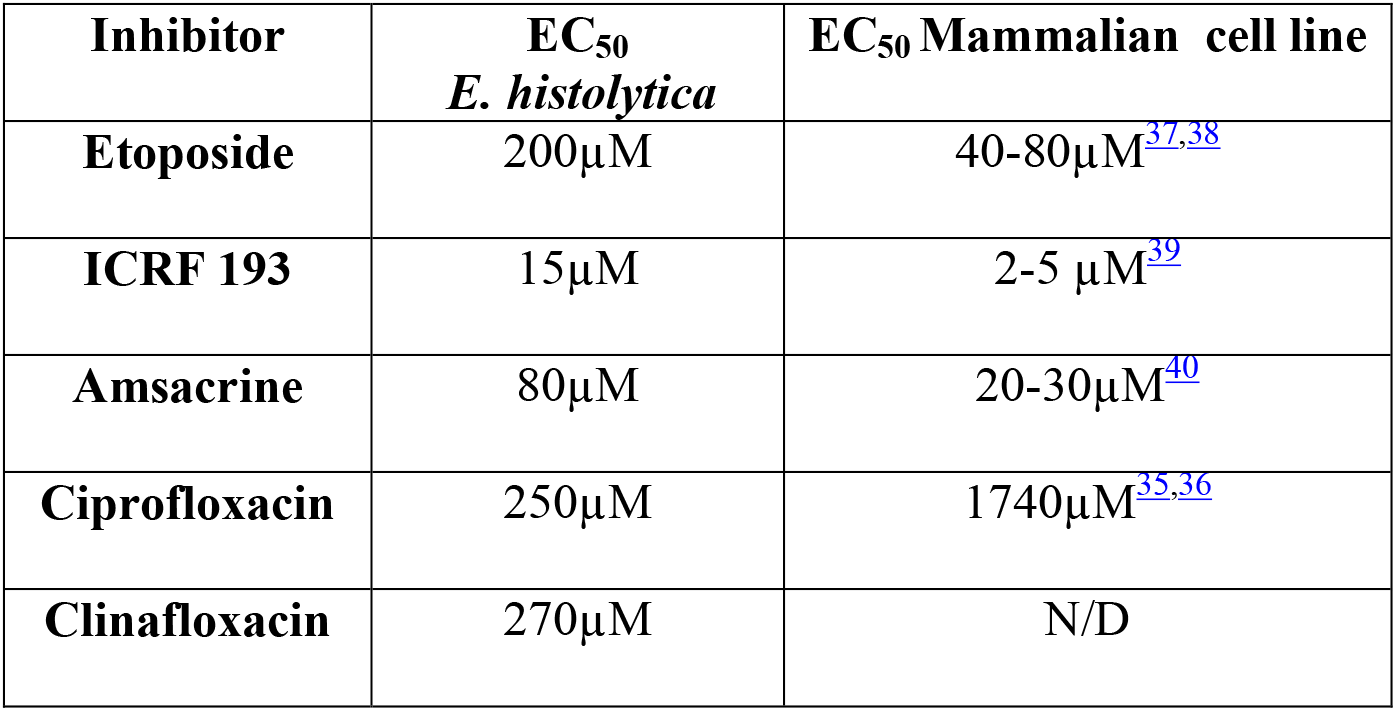
Comparison of EC_50_ of various TopoII inhibitors against *Eh* to that of mammalian cell lines. (N/D: Not determined)

Two crucial regions in hTopo II viz, PLRGKXLNVR (motif I) and Q/MXLMM (motif II) are reported to interact with etoposide [30]. Topo II aminoacid sequence alignment of different *Entamoeba* species with that of hTopo IIα shows that the motif I is mostly conserved. The Q/MXLMM motif on hTopoIIα interacts with etoposide primarily via the M_762_ and M_766_ residues. Mutation of these residues has shown to interfere with the etoposide binding property and increase resistance to this drug [31, 32]. The motif II is mostly conserved among different species of *Entamoeba;* however, it is different from the motif II in hTopoIIα. The key drug interacting methionine residues are absent in all *Entamoeba* species and are naturally substituted with V_723_ and A_727_ in *E. histolytica* (Fig 9).

**Fig 9.**
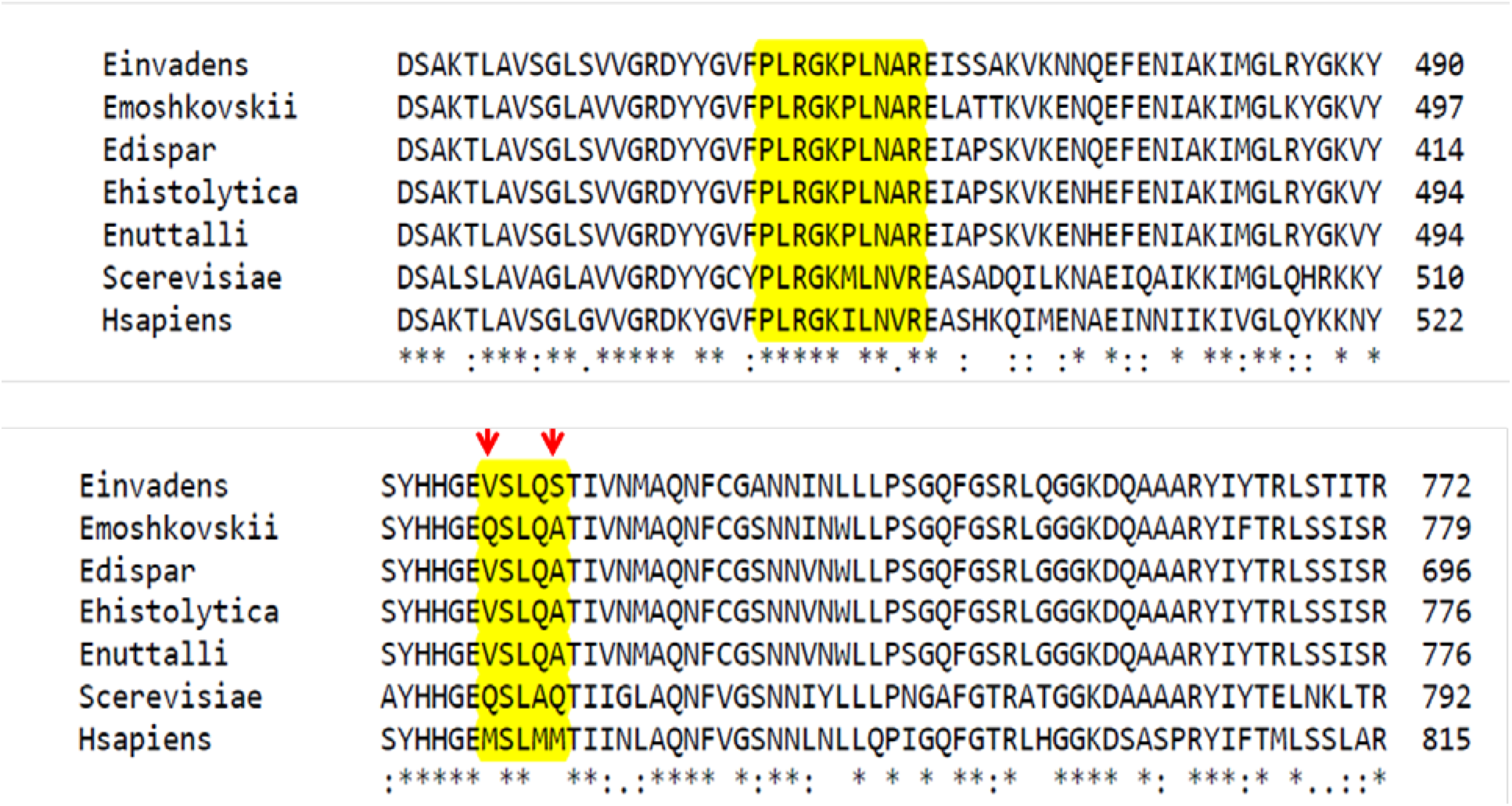
Prediction of etoposide binding motif on *Entamoeba* Topo II. Multiple Sequence Alignment of hTopoIIα and ScTopoII with the ortholog in different *Entamoeba* species showing the PLRGKXLNVR (Motif I) and Q/MXLMM (Motif II) motifs involved in etoposide interaction (highlighted yellow). Most of the interacting residues were conserved between the host and parasite proteins. However, in the motif II, Met_762_ and Met_766_ of hTopoIIα were substituted with Val_723_ and A_727_ in EhTopo II, and these aminoacid replacements were present in all other *Entamoeba* species as well (marked with red arrow). [30]

The drug binding pocket for bisdioxopiperazines like ICRF 193 comprises of 14 residues (7 from each monomer) in eukaryotic Topo II [33, 34]. The histidine residue in this pocket is replaced with glutamine not only in the human parasite but in all other species of *Entamoeba* as well (Fig 10). Alterations in these residues could be the reason for low toxicity of eukaryotic Topo II drugs on *E. histolytica*.

**Fig 10.**
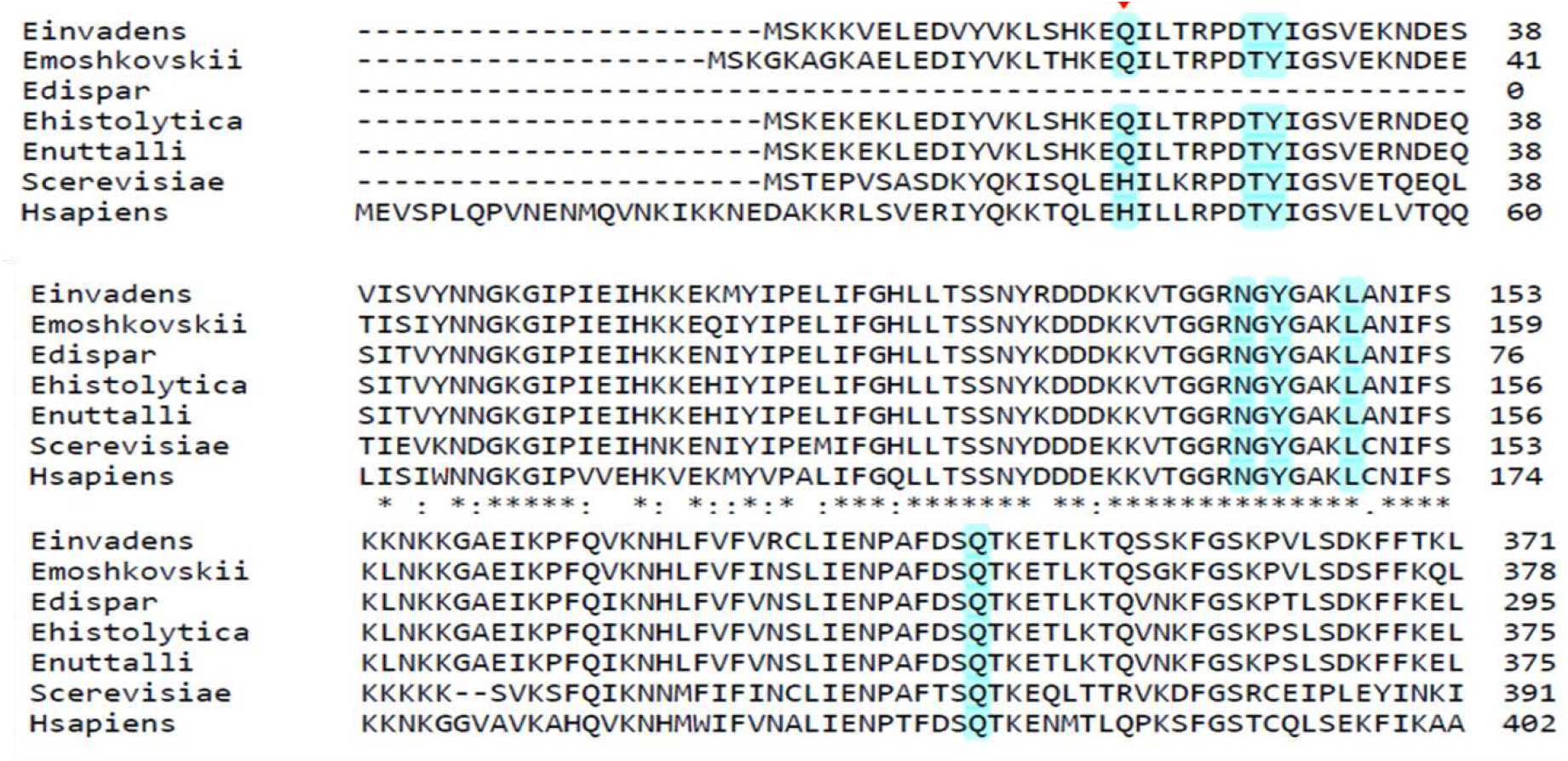
Prediction of ICRF-193 binding motif on *Entamoeba* Topo II. Multiple Sequence Alignment of hTopoIIα and ScTopoII with the ortholog in different *Entamoeba* species showing the residues involved in the interaction with bizdioxopiperazine derivatives like ICRF-193, in eukaryotic Topo II (highlighted blue). The His_41_ involved in the drug binding pocket of hTopoIIα is substituted with Gln_20_ in the EhTopo II. Substitution of His with Gln in the drug binding pocket was conserved in different species of *Entamoeba* (marked with red arrow) [33,34]

### 2.7. Susceptibility of *E. histolytica* to fluoroquinolones targeting prokaryotic Topo IIA

Drugs that target prokaryotic topoisomerase II like DNA gyrases and Topoisomerase IV are primarily fluoroquinolone derivatives. Ciprofloxacin and clinafloxacin are new generation fluoroquinolones with a broad spectrum of antibacterial activity. However, these drugs have a much lower potency against eukaryotic topoisomerase II. Interestingly, both of these drugs were toxic to the survival of *Eh* with an effect comparable to that of etoposide (Fig 11). Further, ciprofloxacin has a profound, direct inhibitory effect on the enzymatic activity of native EhTopo II in comparison to etoposide, affecting the enzyme at a much lower concentration than that of the latter (Fig 12). It is also interesting to note that ciprofloxacin has much lower EC_50_ against *Eh* in comparison to that reported in mammalian cell line [35, 36]. Like in case of *Eh*, ciprofloxacin can reduce the viability of *Ei* and also decrease the encystation efficiency (Fig 13). This opens up scope to further explore newer and better fluoroquinolones in the light of an anti-amoebic drug against this parasite (Table 1).

**Fig 11.**
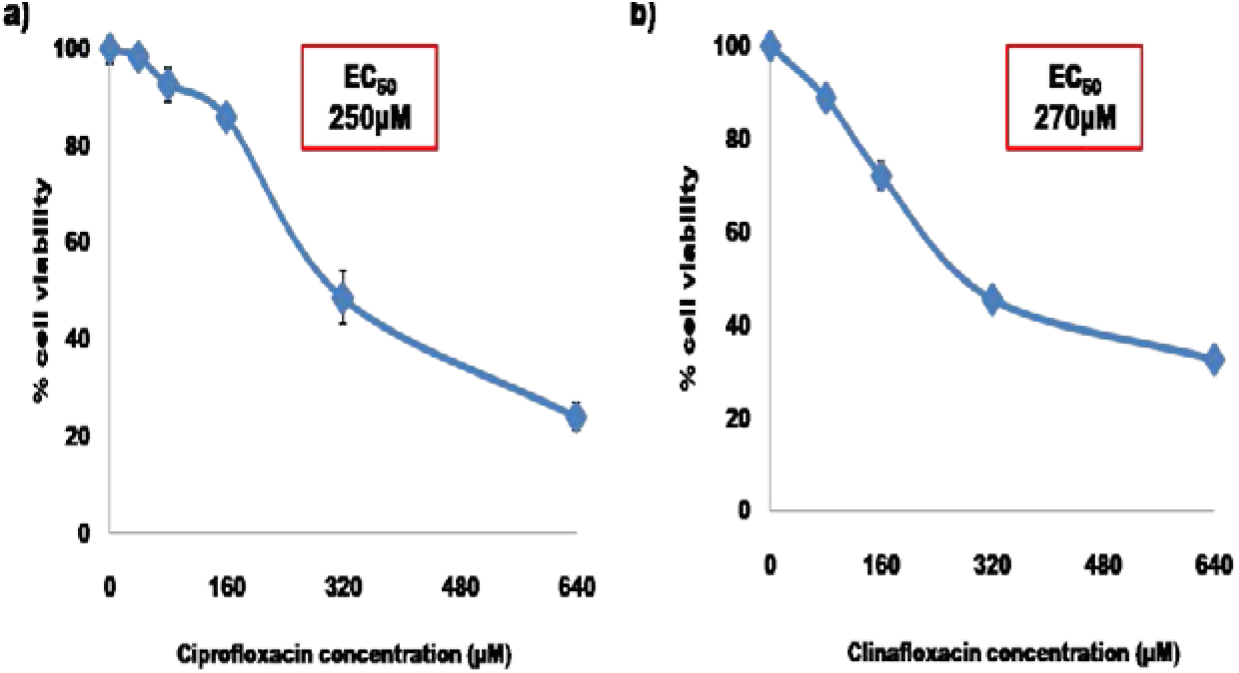
Susceptibility of *E. histolytica* to gyrase inhibitors. Effect of prokaryotic TopoII inhibitors (fluoroquinolones) **a)** ciprofloxacin and **b)** clinafloxacin on the viability of *Eh* post 24h drug treatment.

**Fig 12.**
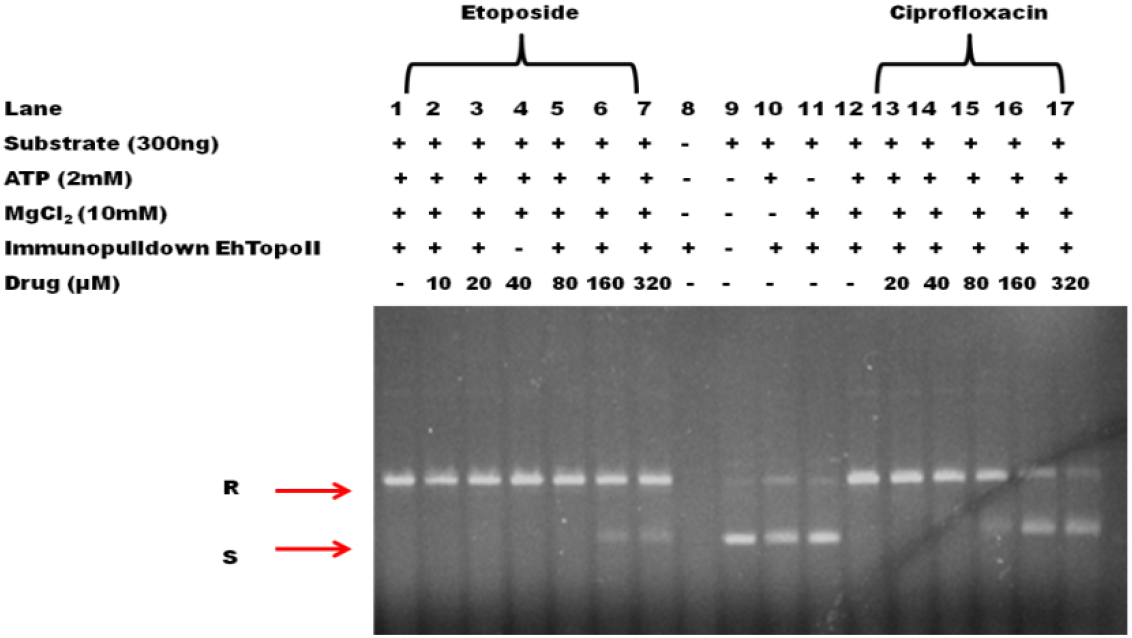
Ciprofloxacin inhibits EhTopo II enzymatic activity. Agarose gel (1%) depicting the inhibitory activity of etoposide and ciprofloxacin on EhTopoII ability to relax negative supercoils. (Lane 1-7: Etoposide treatments; Lane 8-12: different controls; Lane 13-17: Ciprofloxacin treatments.) Ciprofloxacin inhibits the enzymatic reaction at much lower concentrations (160 μM) than the eukaryotic inhibitor, etoposide.

**Fig 13.**
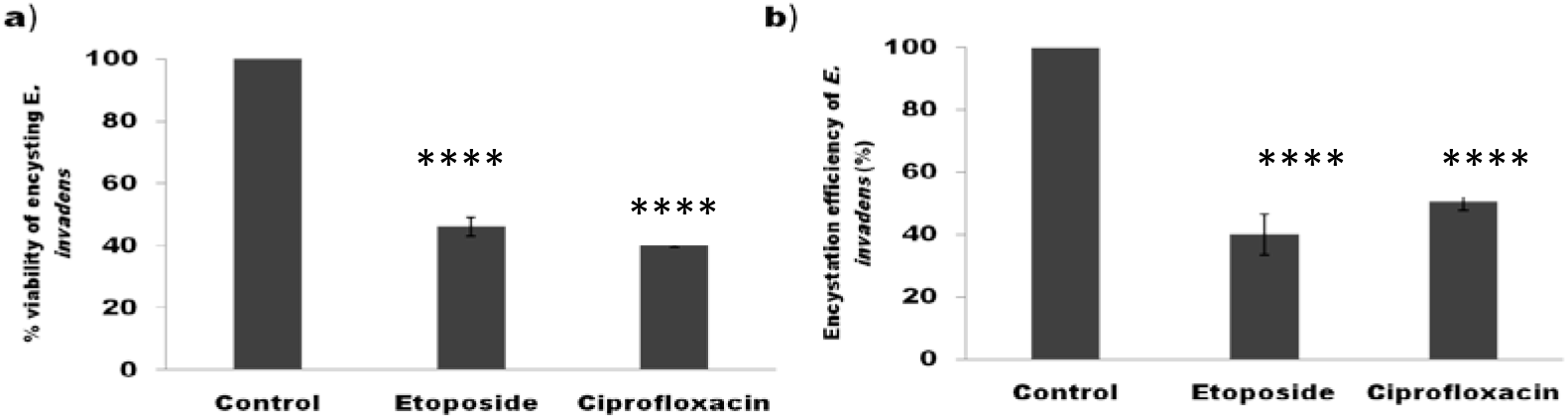
Effect of etoposide and ciprofloxacin on encystation. **a)** The % cell viability and **b)** encystation efficiency of *Ei* after 48h of drug treatment (160μM etoposide and 200μM ciprofloxacin) during encystation. Both treatments reduced the viability as well as the efficiency of encystation (****p<0.00001 w.r.t control)

## 3. Discussion

Topoisomerases, being a crucial player in all DNA processes, is often involved in the stress response and survival of all organisms across the biological world [41, 42]. Like all other organisms, *E. histolytica* also has a pool of genes coding for various types of topoisomerases. The bioinformatic analysis shows that the putative topoisomerases of *E. histolytica* resembled Topo IIIα, Topo IIIβ, Topo II, and SPO11 and are very closely related to their orthologs in *E. invadens*. Although all topoisomerases genes were eukaryotic, (Phylogenetic analysis) recognizable eukaryotic Topo I could not be identified in *Entamoeba*. Among these genes, we observe that topoisomerase II is crucial not only for normal growth and proliferation of the trophozoites but appears to be the most important for the parasites’ response to various stress conditions like heat shock as well as oxidative stress. Similar levels of upregulation of topoisomerase II under these stress conditions are a reported phenomenon in human cancer cells and mediate the excision of chromosomal DNA loops into High Molecular weight (HMW) fragments as a stress response signal under oxidative stress [43, 44].

Stress is one of the most important environmental signals for encystation in *Entamoeba*, leading to the formation of environmentally resistant cysts. As a response to energy deficient environment during glucose starvation and early encystation, trophozoites switch from an actively proliferating stage to a dormant form during which many genes involved in the metabolism and proliferation are downregulated. Likewise, we show that Topoisomerase II, a proliferation marker, also faces a steady decline in the early periods of encystation in *Entamoeba*. Similar stage-specific expression of topoisomerase II has been observed in other parasites as well. For example, in *P. falciparum* the ring stage has a relatively lower level of the protein expression in comparison to trophozoite and schizont stage [18] while in *Leishmania infantum* expression levels varied considerably between infective intracellular amastigotes, proliferative and non-proliferative stages of promastigotes [45]. However, we report that, with the progression of encystation, transcription and translation of topoisomerase II is upregulated in *Entamoeba* around the period of tetranuclear formation, which is one of the key phenomena in a maturing cyst. Further, during this stage of encystation, the enzyme colocalizes on the newly forming nuclei in *E. invadens* suggesting that it may be necessary to remove the topological strains that occur in the DNA during this nuclear event. Also, the significant drop in viability and encystation efficiency upon its downregulation suggests that *Entamoeba* topoisomerase II is indeed important for the proper stage conversion of this parasite.

Over the years, topoisomerase II has emerged as a good drug target and consequently led to the development of an extensive array of medicines as anti-cancer and anti-bacterial agents. As already mentioned, it is extensively explored as a potential drug target in many other parasites as well. We showed that *E. histolytica* has a lower susceptibility to eukaryotic topoisomerase targeting drugs, especially etoposide while fluoroquinolones that target prokaryotic gyrases and topoisomerase IV were significantly toxic to the enzyme activity as well as parasite viability. Sequence analysis of EhTopo II with human TopoIIα show that specific key residues involved in the drug interaction with etoposide and ICRF 193 are naturally substituted in *Eh* and this may be a contributing factor to the poor performance of these drugs towards the parasite in comparison to mammalian cells. So it may be concluded that the eukaryotic TopoII acquired mutation during the evolution that makes it etoposide susceptible. On the other hand, gyrase inhibitors like fluoroquinolones have reported lower potency against higher eukaryotes, including humans [46]. Chemical modifications to the side chains of fluoroquinolones have shown to enhance or decrease its potency towards different organisms [47, 48]. Similar susceptibility to fluoroquinolones, in comparison to mammalian cells, is also reported in case of other parasites like *P. falciparum* and *L. donovanii* and is being explored as a drug target as well [18–21]. Another bacterial gyrase inhibitor GSK299423 (a piperidinylalkylquinoline) has been reported to be extremely effective against *P. falciparum* with 100 times higher potency than against mammalian cell lines [49]. In addition, computational design of newer fluoroquinolones with enhanced potency against parasitic topoisomerase is already gaining traction and thus opens up possibilities for better anti-parasitic drugs [50, 51]. Hence, high potency of fluoroquinolones to *E. histolytica*, low toxicity towards human cells and ease of chemical modification in its side chains are advantages for designing these drugs to specifically target EhTopo II.

Limited treatment methods and potential drug resistance have accelerated the need for newer therapies to tackle amoebiasis. The topoisomerase II of *Eh* is crucial for the stress response and formation of mature cysts and shares only 45% identity with the human counterpart. As we already show that ciprofloxacin is toxic not only to the proliferating trophozoite stage but also severely reduces encystation, designing newer variations of fluoroquinolones that can specifically target *E. histolytica* with minimum side effects to the host could be the answer to not only treat the disease but also prevent the host-to-host transmission of amoebiasis.

## 4. Materials and Methods

### 4.1. In silico analysis

#### 4.1.1. Multiple sequence analysis

Genes annotated as putative topoisomerases from *Eh* and *Ei* were identified from AmoebaDB. Protein sequences of topoisomerases from different organisms were retrieved from the UniProt database (http://www.uniprot.org/) and were aligned with putative topoisomerases of *Entamoeba* using ClustalOmega (http://www.ebi.ac.uk/).

#### 4.1.2. Phylogenetic tree construction

Phylogenetic analysis was performed to understand the categories of topoisomerases the seven in *Entamoeba* fall into. Protein sequence from 50 different topoisomerases covering all known categories, belonging to a range of prokaryotic and eukaryotic organisms like bacteria, fungi, protozoa, plant, and animals were retrieved using UniProt database and aligned with the putative topoisomerases of *Eh* and *Ei* using Clustal W. Using this alignment, phylogenetic tree was constructed with MEGA 7 [52] software by the Maximum Likelihood method and evolutionary distance between sequence pairs was analyzed by the WAG model. Reliability of the model was assessed by bootstrapping with 1000 iterations.

#### 4.1.3. Conserved domain identification and analysis

Conserved motifs on putative *Entamoeba* topoisomerases were identified using MEME software and further analyzed using NCBI-CDS (http://www.ncbi.nlm.nih.gov/Structure/cdd/wrpsb.cgi) [53], and nuclear localization sequences (NLS) was predicted using cNLS mapper (http://nls-mapper.iab.keio.ac.jp/cgi-bin/NLS_Mapper_form.cgi) [54].

### 4.2. Growth, stress induction, and encystation of *Entamoeba*

Trophozoites of *E. histolytica* strain HM-1:1MSS and *E. invadens* IP-1 were grown at 37 °C and 25 °C respectively in Trypticase-Yeast Extract-Iron-Serum (TYI-S-33) medium containing 10% heat-inactivated adult bovine serum, 125μl/100ml streptomycin-penicillin G and 3% vitamin mix [55,56]. As in vitro encystation of *Eh* is not feasible yet, *Ei* is used as the model organism for the same. Late log phase cells were chilled to detach and harvested by centrifugation at 1500rpm for 5min at 4°C. 5×10^5^ cells/ml were transferred to 47% Low Glucose (LG) medium (TYI without glucose diluted 2.12 times with water, 2% heat inactivated adult bovine serum, 2.5% vitamin mix, 125μL/100ml antibiotic). Cysts cultured in LG medium were collected after 12, 24, 36, and 48h.

Cells were subjected to glucose starvation by overnight incubation in TYI-S-33 media devoid of glucose. Trophozoites of *Eh* and *Ei* were incubated at 42°C and 37°C respectively for 1hr for heat shock. Oxidative stress was induced by incubating the cells in media containing 1.0mM of H_2_O_2_ for one hour [57, 58].

### 4.3. RNA isolation, cDNA synthesis, and real-time RT-PCR

The total RNA was isolated using Trizol reagent (Ambion, USA) following standard protocol. The RNA quantity was measured using a NanoDrop spectrophotometer. The isolated RNA were treated with DNase I (Fermentas, USA) at 37°C for 1hr followed by enzyme inactivation at 60°C for 5 min. Genomic DNA contamination was checked by PCR amplification without RT, of actin or ARF gene. Total RNA and genomic DNA were used as template for negative and positive control, respectively.

The cDNA was synthesized from 2μg of purified RNA template in 20 μl reaction volume using OligodT by First Strand cDNA synthesis kit (BioBharati, India) at 42°C for 50 min. The enzyme was deactivated by incubating at 70°C for 15 min. Real-time RT-PCR was carried out following our previously reported standard protocol in Eppendorf Master Cycler RealPlex, using 200ng cDNA, 0.3μM gene specific primers (S1 Table) and PowerUpTM SYBR Green Master Mix (Thermo Fischer Scientific, USA).The fold change of transcript expression was calculated using the ΔΔC_T_ method with respect to housekeeping genes like ADP-ribosylation factor (ARF) or actin. The specificity of the amplicon was validated through melting curve analysis.

### 4.4. Cloning, expression, and purification of truncated EhTopo II

#### 4.4.1. Cloning of *EhTopoII* fragment into a bacterial expression system

Despite using several combinations of vector-strain bacterial expression systems, the full length (4050bp) expression of putative EhTopoII (EHI_120640) was not successful. Hence, a 1127bp long fragment (from 760bp-1887bp) containing no tandem repeats of rare codons was PCR amplified from *Eh* genomic DNA with the specific primers EhTopoIIfgS*Bam*HI and EhTopoIIfgAS*Xho*I (S1 Table) using High Fidelity Taq polymerase (Thermo Fischer Scientific, USA) and subcloned into pGEMT-Easy (Promega, USA) via TA ligation. This 1127bp fragment shared 77% DNA sequence identity with the ortholog in *Ei* (EIN_145900) with no gaps and carried regions of the putative catalytic site. Positive clones identified by blue-white screening and confirmed by *Bam*HI-*Xho*I double digestion were sequenced and further cloned into the pET21a expression vector. Restriction digestion was done for clone confirmation, and recombinant vector was transformed into *E. coli* BL21 (DE3) expression strain.

#### 4.4.2. Expression and purification of recombinant truncated EhTopo II protein

Overnight cultures of *E. coli* BL21(DE3) strain carrying recombinant pET21a vectors were diluted to OD_600_ 0.05 and grown at 37°C till the OD_600_ was 0.6-0.8 at which recombinant protein expression was induced using 1mM IPTG for 4 hours at 37°C. The recombinant protein was solubilized using 0.25% S-lauryl sarcosine and further purified using Ni-NTA affinity chromatography. The purified recombinant protein fragment was resolved on 12% SDS-PAGE.

### 4.5. Generation and purification of Anti-EhTopo II polyclonal antibody

Polyclonal antibody against recombinant EhTopo II fragment was commercially raised in rabbits (Abgenex, India) and purified from the crude sera using Protein-A sepharose (Invitrogen, USA) affinity chromatography. The sera were loaded over a pre-equilibrated Protein-A sepharose column following 1:1 v/v dilution in PBS (pH 8). After sufficient washing to remove all unbound proteins, the antibody was eluted using 100mM glycine (pH 2.8), and at least 20 fractions of 1mL each were collected into tubes containing 100μL of 1M Tris (pH 9). The absorbance of the fractions at 280nm was measured, and the antibody-containing samples were pooled, dialyzed over several changes of 1x PBS and concentrated.

### 4.6. Total protein extraction from *Entamoeba*

The *Entamoeba* cells grown at different conditions were harvested, washed with 1x PBS and resuspended in lysis buffer containing 20mM Tris (pH 7.5), 1mM EDTA, 200mM NaCl, 1mM PMSF, 1μg/mL of leupeptin and pepstatin,0.3μM of aprotinin, 15μM E-64, 1% Triton-X and 0.1% SDS and incubated for 30mins. The samples were then sonicated, centrifuged for 10mins at 10,000 rpm at 4°C, and the supernatant was collected for further experiments. This method was followed for samples from all conditions, including different kinds of stress and different hours of encystation.

### 4.7. Western blot analysis

Recombinant as well as native protein samples were run on 12% SDS-PAGE and blotted onto PVDF membrane at 70V for 2h. The membrane was blocked for 2h using 3% BSA in PBST (0.1% Tween 20 in 1xPBS) and probed with anti-EhTopo II antibody (1:2000) for 2h at 4°C. Further, it was washed extensively with PBST and incubated with HRP conjugated goat anti-rabbit IgG antibody (1:5000) for 90min at room temperature. After washing thrice with PBST, the membrane was developed using Luminata Classico Western HRP substrate (Merck Millipore, USA) and detected in ImageQuant LAS500 imager (GE Life Science, USA).

### 2.8. Nuclear extract preparation from *Entamoeba*

The harvested log phase trophozoites were washed thrice with 1x PBS and resuspended in 5x volume ice-cold TEMP buffer (10 mM Tris-HCl, pH 7.5, 1 mM EDTA, 4 mM MgCl2, 0.5 mM PMSF), containing 2 μg/ml leupeptin and aprotenin, 1μg/ml pepstatin and μM E-64. After 30 min incubation on ice, the cells are lyzed using 0.5% Triton-X treatment for 10 mins. The lysate is overlaid on a 1.5M sucrose cushion and spun at 12,000xg for 10mins. The pure nuclei pellet so obtained is resuspended in TEP buffer (10 mM Tris-HCl, pH 7.5, 1 mM EDTA, 0.5 mM PMSF) containing 0.35M NaCl and all protease inhibitors and incubated on ice for 45 mins followed by centrifugation at 12,000xg for 15 min. The supernatant is the nuclear extract which was assayed for topoisomerase II activity.

### 4.9. Immunoprecipitation of native Topoisomerase II

Functional characterization of *Entamoeba* Topoisomerase II was carried out by immunoprecipitation of the protein from crude nuclear extracts of log phase trophozoites using anti-EhTopo II antibody. The nuclear extract was pre-cleared with pre-immune rabbit IgG (10 μg/ml) and 20 μl of protein-agarose beads (BioBharati, India) 1h at 4°C. The pre-cleared supernatant was collected by centrifugation and incubated with purified anti-EhTopo II IgG (10μg/ml) at4 °C following which, 20 μl of protein-A agarose beads were added to precipitate the antigen-antibody complex and incubated at 4°C for 2h. Agarose beads with bound immune complexes were then harvested by centrifugation at 4°C and washed with nuclear extract buffer. The immunoprecipitated topoisomerase II was confirmed by Western blotting and used for further assays.

### 4.10. Topoisomerase II assay

The relaxation assay was carried out according to the protocol reported by Chakraborty and Majumder, 1987 [59]. The standard relaxation assay mixture contained: 25mM Tris-HCl, pH 7.5, 10mM MgCl_2_, 0.1mM EDTA, 1mM DTT, 2mM ATP, 50mM NaCl, 10% glycerol, 500ng of supercoiled pure pL4440 plasmid substrate and different concentrations of nuclear extract or immunoprecipitated protein solution as enzyme source. The reactions were carried out for 30min at 25°C and 37°C for extracts from *Ei* and *Eh* respectively and arrested by adding dye containing 1% SDS and 10mM EDTA. The samples were electrophorized on 1% agarose gel at 1V/cm overnight at room temperature and stained with ethidium bromide.

### 4.11. Staining and confocal microscopy

*Entamoeba* cells were harvested, washed with PBS (pH 7.6) and fixed with 2% p-formaldehyde for 30mins at room temperature. The fixed cells were washed thoroughly and permeabilized with 0.1% Triton-X for 5mins blocked for 1h with 3% BSA and incubated overnight at 4°C with anti-EhTopo II antibody (1:100) in 1xPBS containing 1% BSA. It was followed with further washes and incubation at room temperature with TRITC conjugated anti-rabbit IgG (1:400) (Sigma, USA) for 1h. The nucleus was stained using 10μg/mL DAPI. The stained cells were visualized and analyzed using Olympus FluoView FV1000 confocal microscope and software.

### 4.12. dsRNA mediated RNA interference studies

#### 4.12.1. Cloning and expression of *EhTopoII* and *EiTopoII* specific dsRNA

In order to achieve downregulation at the RNA level, a 300bp and 250bp fragment within *EhTopoII* and *EiTopoII*, respectively were selected as targets for RNA interference. Specificity of the dsRNA was ensured by the selected regions within *Entamoeba TopoII* that did not contain 19mer homology to any other genes in *Eh* and *Ei*. These regions were amplified using specific primers EiTopoIIdsF*Hin*dIII-EiTopoIIdsR*Xho*I and EhTopoIIdsF*Xba*I-EhTopoIIdsR*Xho*I (S1 Table), subcloned into pGEMT-Easy TA vector and then into expression vector pL4440, a gift from Andrew Fire (Addgene) (http://n2t.net/addgene:1654) which is flanked by T7 promoter on either side of its MCS. Recombinant plasmids carrying the inserts were transformed into the RNase III-deficient, *E. coli* HT115 cells. The dsRNA expression was induced by 1mM IPTG when OD_600_ reached 0.6, followed by incubation at 37°C for 4h.

#### 4.12.2. Extraction and purification of dsRNA

The gene specific dsRNA was extracted using water saturated phenol: chloroform: isoamyl alcohol (25:24:1) followed by phase separation at high speed centrifugation. The RNA from the aqueous layer was precipitated with equal volumes of isopropanol, further washed with 70% ethanol, air dried and resuspended in nuclease-free water. Single-stranded RNA and DNA contaminants were eliminated by treating the sample with 0.2μg/μL RNaseA (Sigma, USA) and 0.1U/μL DNase (Thermo Fischer Scientific, USA) for 1h at 37°C. The samples were further purified using Trizol, following standard protocol. The dsRNA from a 150 bp region in the flavin mononucleotide based fluorescence protein of *B.subtilis* (BS1), with no sequence similarity to *Entamoeba* genome, was used as negative control dsRNA.

#### 4.12.3. dsRNA mediated downregulation of *Entamoeba* by soaking

*Eh* and *Ei* trophozoites were incubated in TYI or LG media containing 200 μg/ml of the respective, purified TopoII-specific dsRNA for various time points, based on the experiment, and the silencing efficiency was calculated using real-time RT-PCR [29]. All RNAi studies were carried out with two controls: one without dsRNA and one with non-specific dsRNA (from *B. subtilis*, as mentioned above).

### 4.13. Determination of cell viability and encystation efficiency

Effect of *Entamoeba Topo II* silencing and potency of various topoisomerase II drugs was studied by assessing their effect on viability. Cell viability was determined by trypan blue dye exclusion assay. Harvested cells, following various treatments, are resuspended in 1x PBS and mixed with equal volumes of 0.4% trypan blue (Himedia, India). Dead cells take up the dye and appear blue, while live ones remain white. The cell count is calculated at the start of the experiment as well as following dye treatment, using a Neubauer hemocytometer.

After 48h of encystation, the total cell number (cyst+trophozoite) was calculated, and the harvested cells were treated with 0.5% sarcosine for 10mins to lyze the trophozoites. The number of cysts was then calculated and the encystation efficiency is represented as (cyst x 100)/(cyst+trophozoite).

### 4.14. Structure modeling and analysis

The three-dimensional protein structure of EhTopo II was predicted by homology modeling using SWISS-Model server (https://swissmodel.expasy.org/). The template structure was obtained from PDB based on sequence similarity and, refined using ModRefiner server (http://zhanglab.ccmb.med.umich.edu/ModRefiner/). The generated structure was validated by the Ramachandran plot using PROCHECK. PyMol software was used for the visualization of the protein structure.

## Abbreviations

*Eh*: *E. histolytica*
*Ei*: *E. invadens*
EhTopo II: *E. histolytica* topoisomerase II
EiTopo II: *E. invadens* topoisomerase II
SPO11: Meiotic recombination protein
Topo: Topoisomerase

## Acknowledgement

SKG would like to thank DBT, Govt. of India for partial funding of this project. The authors thank Indian Institute of Technology Kharagpur for the research facilities and Fellowship to SSV. The authors like to acknowledge FIST, DST, Govt. of India for the confocal facility. The *E. coli* HT115 (DE3) strain was a kind gift from the Caenorhabditis Genetics Centre, funded by the NIH National Centre for Research Resources, USA.

## Supporting information caption

**Tables**

S1 Table: List of primers

S2 Table: List of putative topoisomerase in *Entamoeba*

**Figures**

S1 Fig: Comparison of FPKM values of different topoismerases of *E. invadens* during encystation

S2 Fig: Expression and purification of recombinant truncated EhTopo II.

S3 Fig: Western blot analysis of native and recombinant Topo II

S4 Fig: Cloning and expression of *Entamoeba* Topo II specific dsRNA and dsRNA mediated silencing of TopoII in *Entamoeba*.

