## supporting information figures for "Stress-responsive *Entamoeba* topoisomerase II: a potential anti-amoebic target"

### S1 Fig. Comparison of FPKM values of different topoismerases of *E. invadens* during encystation

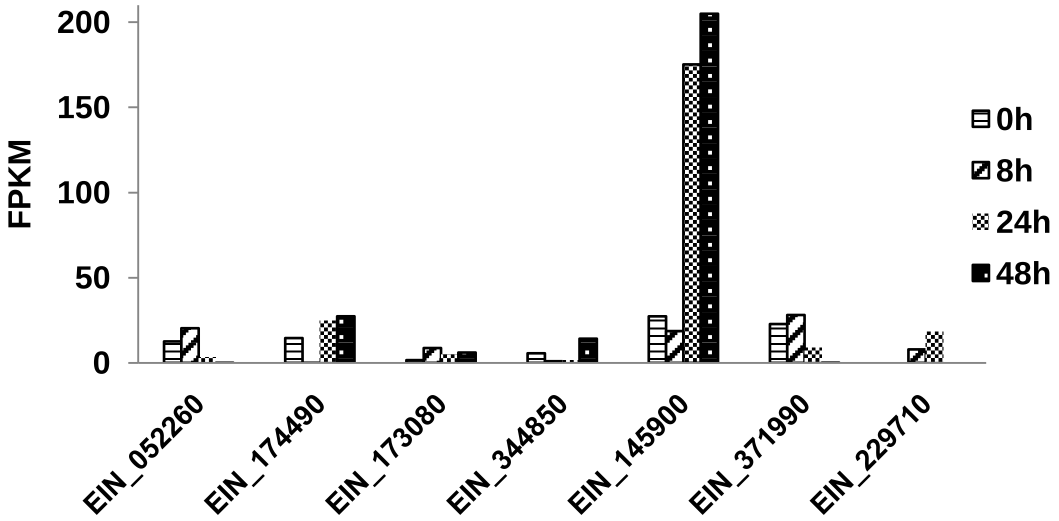


**S1 Fig. Comparison of FPKM values of different topoismerases**. Microarray data of putative topoisomerases from *E. invadens* shows that EIN_145900, which is a putative Topoisomerase II, was highly upregulated during encystation. ( Data were obtained from AmoebaDB*)


* Jeelani G, Sato D, Husain A, Cadiz AE & Sugimoto M (2012) Metabolic profiling of the protozoan parasite *Entamoeba invadens* revealed activation of unpredicted pathway during encystation. *PLoS ONE*. 7(5), e37740.

### S2 Fig. Expression and purification of recombinant truncated EhTopoII

c)

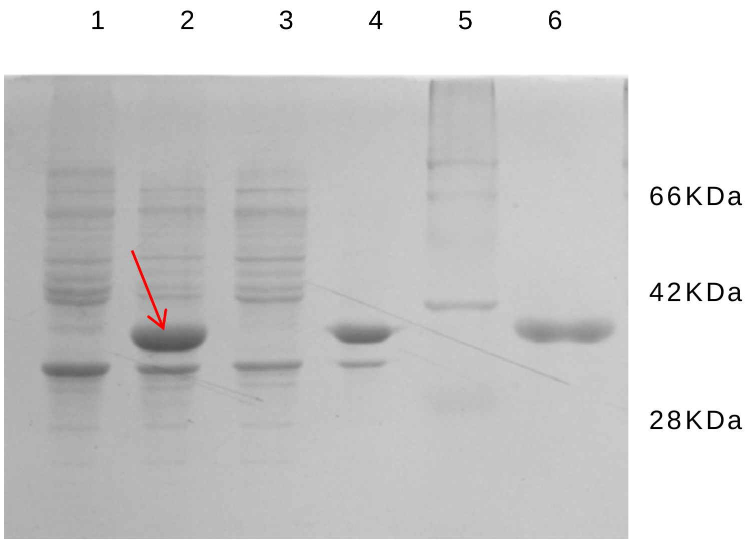


**S2 Fig. Expression and purification of recombinant truncated EhTopo II.**12% SDS PAGE gel showing expression of truncated insoluble EhTopoII protein in BL21(DE3) strain upon 0.8mM IPTG induction. **Lane 1:** Uninduced BL21(DE3) **Lane 2:** Induced BL21(DE3) **Lane3:** Induced pellet fraction **Lane 4:** Induced supernatent fraction **Lane 5:** Protein ladder **Lane 6:** Recombinant TopoII solubilized from inclusion bodies (6M urea) and purified using Ni-NTA affinity chromatography

### S3 Fig. Western blot analysis of native and recombinant Topo II

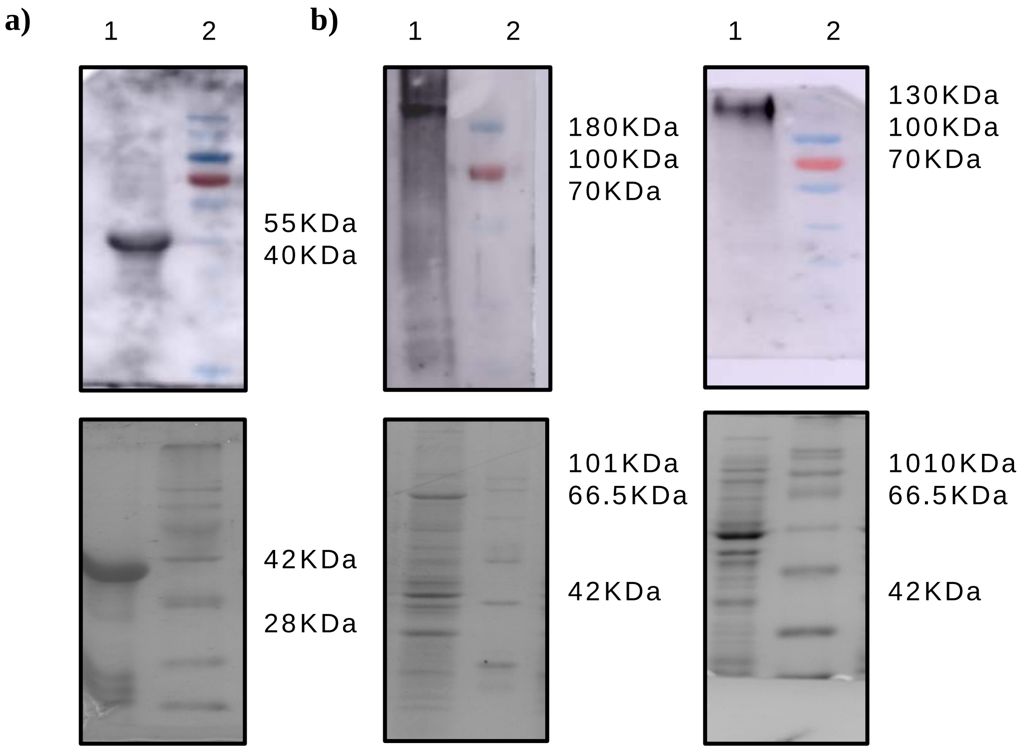


**S3 Fig. Western blot analysis assaying anti-Topo II antibody** against **a)** recombinant EhTopoII fragment and total cell lysate from **b)** *E. invadens* and **c)** *E. histolytica*

### S4 Fig. Cloning and expression of *Entamoeba* Topo II specific dsRNA and dsRNA mediated silencing of TopoII in *Entamoeba*

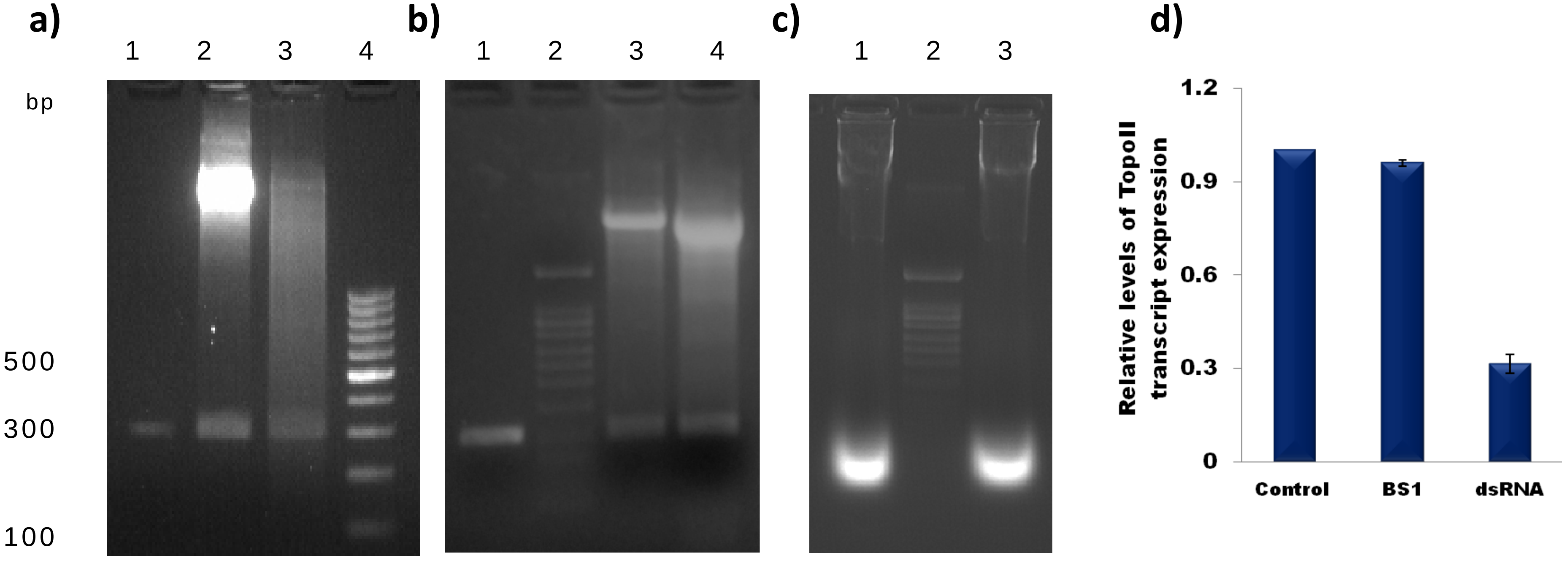


**S4 Fig.** Agarose gel showing PCR amplification of **a)** EiTopoIIds and **b)** EhTopoIIds gene fragment and positive clones in pGEMT_Easy and pL4440 vectors respectively. **c)** Purified EhTopoII and EiTopoII dsRNA extracted from recombinant HTII5 **d)** Relative Topo II transcript expression (calculated relative of ARF) after 48h of encystation in *E. invadens* during dsRNA mediated gene silencing. BS1-non specific dsRNA from *Bacillus subtilis* gene
