## supporting information tables for "Stress-responsive *Entamoeba* topoisomerase II: a potential anti-amoebic target"

**S1 Table: List of primers**

| S.No. | Primer Name | Sequence (5'-3') |
|---|---|---|
| Primers for RT-PCR analysis | | |
| 1 | Eh125320F | C A G G A A C A T C T A A A C C G A G T G A |
| | Eh125320 R | G T G T T G A A T G A C T G A G A C A A C T C G |
| 2 | Eh194510F | ATTACTGCATGTGGCTTTC |
| | Eh 194510R | CATAACAAACCTTCTTCTGAT |
| 3 | Eh120640F | CTG CAT GTG GCT TTC CTG |
| | Eh120640R | CCT TCT TCT GAT CAA TTA AGG AAA G |
| 4 | Eh038920F | TGGAATTGGTACTGATGCAA |
| | Eh 038920R | CTCTTGTTGAATTGTGCCC |
| 5 | Eh042880F | GAA TAA GTG TTG GTG TTG AAT GTT C |
| | Eh 042880R | CAA CTC TTC GGG TAA TGA AAT |
| 6 | Ei371990F | CGGTGTAATTCCTTGTTGTTTGGTT |
| | Ei371990R | CCCGCTTTGTGAGACTCGAC |
| 7 | Ei229710F | GCGTCTTTGTCAGTACTATCTCA |
| | Ei229710R | CACCACAACATCACCCAACTTTT |
| 8 | Ei145900F | AAGAAAGACGACGACACAAGC |
| | Ei145900R | TCTTCGTCTTCATCCGAATTAC |
| 9 | Ei052260F | CGCGAGGGGGTTTGAAACG |
| | Ei052260R | CGACGTCTCTCCCATTTCTTGTG |
| 10 | Ei174490F | GTGTGTGGTGGGGAGATGTG |
| | Ei174490R | GCACTTCTCACAAGGAACACCA |
| Cloning Primers | | |
| 11 | EhTopoIIfgS*Bam*HI | GGATCCATGTATGTTGGTGATGAACCAATAGT |
| | EhtopoIIfgAS*Xho*I | CTCGAGATCTGAACCAGCTCCATCATATG |
| 12 | EhTopoIIdsF*Xba*I | TCTAGAATAAGAAGAGCATATGATATTGC |
| | EhTopoIIdsR*Xho*I | CCGCTCGAGGTCTACATGAGTTCCTCCATG |
| 13 | EiTopoIIdsF*Hind*III | AAGCTTTTGCAATCGACCATTGTCA |
| | EiTopoIIdsR*Xho*I | CTCGAGATCGGTAGACCATCCCGTT |

### S2 Table: List of putative topoisomerases in *Entamoeba*

| Putative topoisomerase in *E. histolytica* | Protein length | NLS (aminoacid residues) | TM helix Sequence | Identical protein in *H. sapiens* (% identity) | Ortholog in *E. invadens* |
|---|---|---|---|---|---|
| EHI_038920 | 607 | 342-408 | 201-221 | Topo IIIα (44% ) | EIN_052260 (68%) |
| EHI_042880 | 840 | 594-623 | 718-740 | TopoIIIβ (38%) | EIN_174490 (64%) |
| EHI_087330 | 400 | 263-293 | | PAPD5 isoform2 (30% ) | EIN_173080 (50%) |
| EHI_073170 | 365 | 58-88 | 131-155<br>214-235 | PAPD5 Isoform2 (27%) | EIN_344850 (64%) |
| EHI_120640 | 1349 | 1175-1270 | --- | TopoIIα (45%) | EIN_145900 (72%) |
| EHI_125320 | 340 | 294-324 | --- | SPO11 (26%) | EIN_371990 (48%) |
| EHI_194510 | 293 | 83-114 | --- | SPO11 (27%) | EIN_229710 (23%) |
